# Genomic mosaicism reveals developmental organization of trunk neural crest-derived ganglia

**DOI:** 10.1101/2024.09.25.615004

**Authors:** Keng Ioi Vong, Yanina D. Alvarez, Geoffroy Noel, Scott T. Barton, Changuk Chung, Robyn Howarth, Naomi Meave, Qingquan Zhang, Fiza Jiwani, Chelsea Barrows, Arzoo Patel, Jiang Xiong Wang, Neil Chi, Stephen F. Kingsmore, Melanie D. White, Xiaoxu Yang, Joseph G. Gleeson

**Author notes:** These authors contributed equally (Keng Ioi Vong, Xiaoxu Yang).

## Abstract

The neural crest generates numerous cell types, but conflicting results leave developmental origins unresolved. Here using somatic mosaic variants as cellular barcodes, we infer embryonic clonal dynamics of trunk neural crest, focusing on the sensory and sympathetic ganglia. From three independent adult neurotypical human donors, we identified 1,278 mosaic variants using deep whole-genome sequencing, then profiled allelic fractions in 187 anatomically dissected ganglia. We found a massive rostrocaudal spread of progenitor clones specific to sensory or sympathetic ganglia, which unlike in the brain, showed robust bilateral distributions. Computational modeling suggested neural crest progenitor fate specification preceded delamination from neural tube. Single-cell multiomic analysis suggested both neurons and glia contributed to the rostrocaudal clonal organization. CRISPR barcoding in mice and live imaging in quail embryos confirmed these clonal dynamics across multiple somite levels. Our findings reveal an evolutionarily conserved clonal spread of cells populating peripheral neural ganglia.

**Highlights:**

- Genetic mosaicism and real-time imaging reveal trunk neural crest cellular dynamics.
- DRG or SG cells from different axial levels are more lineage-related than from the same level.
- Cell fate specification of trunk neural crest progenitors occurs before neural tube delamination.
- These aspects of clonal organization are evolutionarily conserved across mammals and avians.

## Introduction

The neural crest (NC) gives rise to diverse ectomesenchymal cell types that help define features of the vertebrate body plan, including the ganglia of the peripheral nervous system, craniofacial skeleton, and smooth muscle, among others. The broad spectrum of NC potential fates has sparked interest in discovering principles underlying cell fate diversification, representing one of the most longstanding and pivotal questions in developmental biology^1,2^.

The early emergence of NC during gastrulation, their highly motile nature, and rapid expansion present significant technical hurdles for precise mapping of NC progenitor clonal dynamics and organization. Conventional lineage tracing of NC has relied on cellular tracers such as radioisotopes or lipophilic dye in avian or amphibian embryos^3-5^. More recently, transgenic technology has made clonal tracing in mice possible. When analyzing a single somite segment with the *Confetti* reporter, it was suggested delaminated trunk NC retained multipotency to generate both the dorsal root ganglia (DRG) and sympathetic ganglia (SG)^6^. Yet, the limited clonal configuration of fluorescent protein combinations precludes a more comprehensive assessment of clonal relationships.

NC developmental fate is thought to be closely coupled with migration paths. Following convergence of neural folds at the dorsal midline, NC cells undergo epithelial-mesenchymal transition, delaminating from the neural tube (NT), to adopt one of two stereotypical migration paths: the ventromedial route through the rostral sclerotome, generating DRG and SG, or the dorsolateral route between the epidermis and dermamyotome, generating melanocytes^7,8^. However, NC induction occurs at a much earlier time point in the medial epiblast, even before neurulation begins^9^. The cellular dynamics between initial NC emergence and delamination remain largely elusive, especially in mammals. Anomalies during NC development cause common developmental malformations with diverse manifestations, collectively known as neurocristopathies and classified based on NC axial origins^10^.

Recent advances in mosaicism analysis offer opportunities to infer clonal dynamics and resolve developmental lineage in a comprehensive manner. During nearly each cell cycle, postzygotic cells acquire somatic mutations that are faithfully inherited by daughter cells, serving as naturally occurring inherent cellular barcode tracers^11,12^. Rapid proliferation during embryogenesis creates clones carrying barcodes, that when assessed in cellular pools, reveal varying allelic fractions (AFs). Leveraging clonal mosaicism within an organism can help reconstruct lineage relationships based on clonal similarities. This approach, termed “mosaic variant barcode analysis (MVBA)”, has proven powerful in deconvolving the lineages in the brain and hematopoietic system^13-15^.

Here, by applying MVBA to three independent human donors, we provide an unbiased, large-scale assessment of the lineage relationships of trunk NC derivatives spanning from cervical to lumbar levels. The evaluation of more than 1,200 clones demonstrates long-range, bilateral spread of DRG or SG-enriched clones across multiple axial levels prior to NT delamination. Clonal analysis indicates ganglia from different axial levels share a closely related lineage, while the DRG and SG located proximally are typically phylogenetically distinct. These lineage relationships are recapitulated in mice by CRISPR barcoding tracing. Furthermore, real-time whole embryo imaging of quail embryos uncovers previously undescribed rostrocaudal, midline-traversing migration routes of NC progenitors. Finally, mathematical modeling of clonal dynamics reveals cell fate restriction before NC delamination as the prevailing mechanism of trunk NC lineage diversification. Thus, by combining MVBA with live imaging, our study uncovers a previously unrecognized developmental organization of trunk NC progenitors that is evolutionarily conserved across vertebrates.

## Results

### Identification of mosaic variants in the dorsal root ganglia and sympathetic chain ganglia

To understand clonal relationships of the sensory and sympathetic ganglia across the body axes, we isolated 94 DRGs and 93 SGs across cervical, thoracic, and lumbar levels from two male (ID06, ID07) and one female (ID08) adult human donors (Figures 1A, S1A). To define the landscape of somatic mosaic variants (MVs), we also collected peripheral organs including the brain, heart, liver, both kidneys, and skin punches. A subset of ganglia and tissue punches from organs were subjected to 30x and 300x whole-genome sequencing respectively, followed by processing with state-of-the-art MV calling and filtering pipeline to identify candidate MVs (Figure S1B, STAR Methods). Next, we conducted ultra-deep massive parallel amplicon sequencing (MPAS), and quantified AFs of more than 1200 bona fide MVs (>500×) for all collected ganglia. (Figures 1B-1D, S1C-S1D). Single-nucleus RNA sequencing of ganglia from ID08 suggested the trunk NC derivatives, including neurons, Schwann cells, and satellite glia cells, together accounted for approximately 80% of sequenced cells in DRG or SG (Figure S1E). Thus, the majority of clonal MVs originated from the trunk NC lineage. Notably, more than 30% of MVs were enriched by at least 1.5-fold in either DRGs or SGs of donors ID06 and ID07 (30.8% and 37.4% respectively, STAR Methods). This proportion was less in ID08, likely because fewer DRGs were isolated due to variable sample quality.

**Figure 1.**
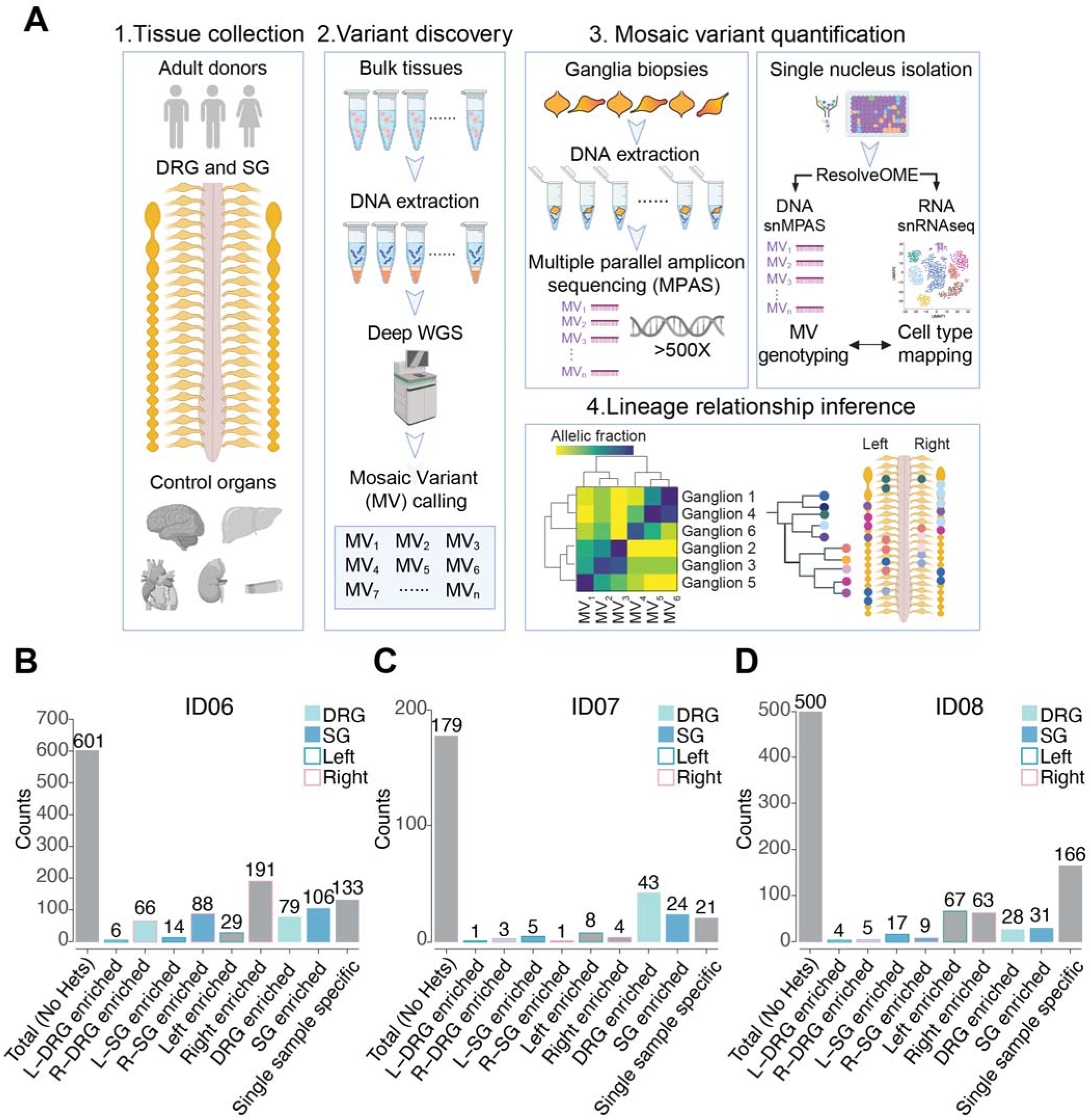
Identification of 1280 bona fide somatic mosaic variants in human DRG and SG from 3 neurotypical control donors. (A) Strategy to deconvolve lineage of sensory (DRG) and sympathetic ganglia (SG): (1) Tissue collection: DRG and SG dissected from 2 male and 1 female donors. Other major organs were also collected to infer clonal relationship; (2) Variant discovery: 300x and 30x whole-genome sequencing (WGS) of bulk peripheral organs and ganglia biopsies respectively, followed by best-practice mosaic variant (MV) calling pipelines, identified candidate MVs. (3) Candidate MVs quantified in dissected tissues or single nuclei isolated from individual ganglia by multiple parallel amplicon sequencing (MPAS) or single-nucleus MPAS (snMPAS) respectively. (4) Lineage tree inference: Variant allelic fractions of validated MVs in individual ganglia analyzed and anatomically mapped. Lineage relationships of ganglia deconvolved by computing clonal similarities between samples and statistical modeling of clonal dynamics. DRG, dorsal root ganglia; SG, sympathetic ganglia. (B-D) Mosaic variant counts identified from donors ID06 (B), ID07 (C), and ID08 (D), classified by tissue and anatomical distribution. Variant detected 1.5x more frequent in a group is defined as enriched. See STAR methods for mathematical quantification. Heterozygous variants (Hets) were excluded. L, left; R, right.

### Rostrocaudal but not dorsoventral origin of trunk neural crest progenitor clones

To reveal distribution of clonal variants, we mapped the AF of each MV onto a schematic body plan (Figures 2A-D, S2A-S2L), which we referred to as a ‘geoclone’. As widely perceived, DRGs and SGs shared variants with spinal cord for ID06 (for which spinal cord was also profiled), consistent with the model in which the neural crest (NC) delaminates from the NT. However, according to the model, MVs were expected to be shared between DRGs and SGs from adjacent spinal levels and potentially unilateral. To our surprise, AF distribution of MVs was often negatively correlated between DRGs or SGs of the same level but was positively correlated between DRGs of different levels or between SGs of different levels. Moreover, there were multiple MVs that spanned either DRGs or SGs up to 12 spinal levels apart but were not locally shared between the DRGs and SGs at the same level. For example, in donor ID07, the MV chr7:47802159-T-A was detected in almost all thoracic SGs but barely detected in DRGs, except at the rostral-most T1 and T2 levels (Figure 2C).

**Figure 2.**
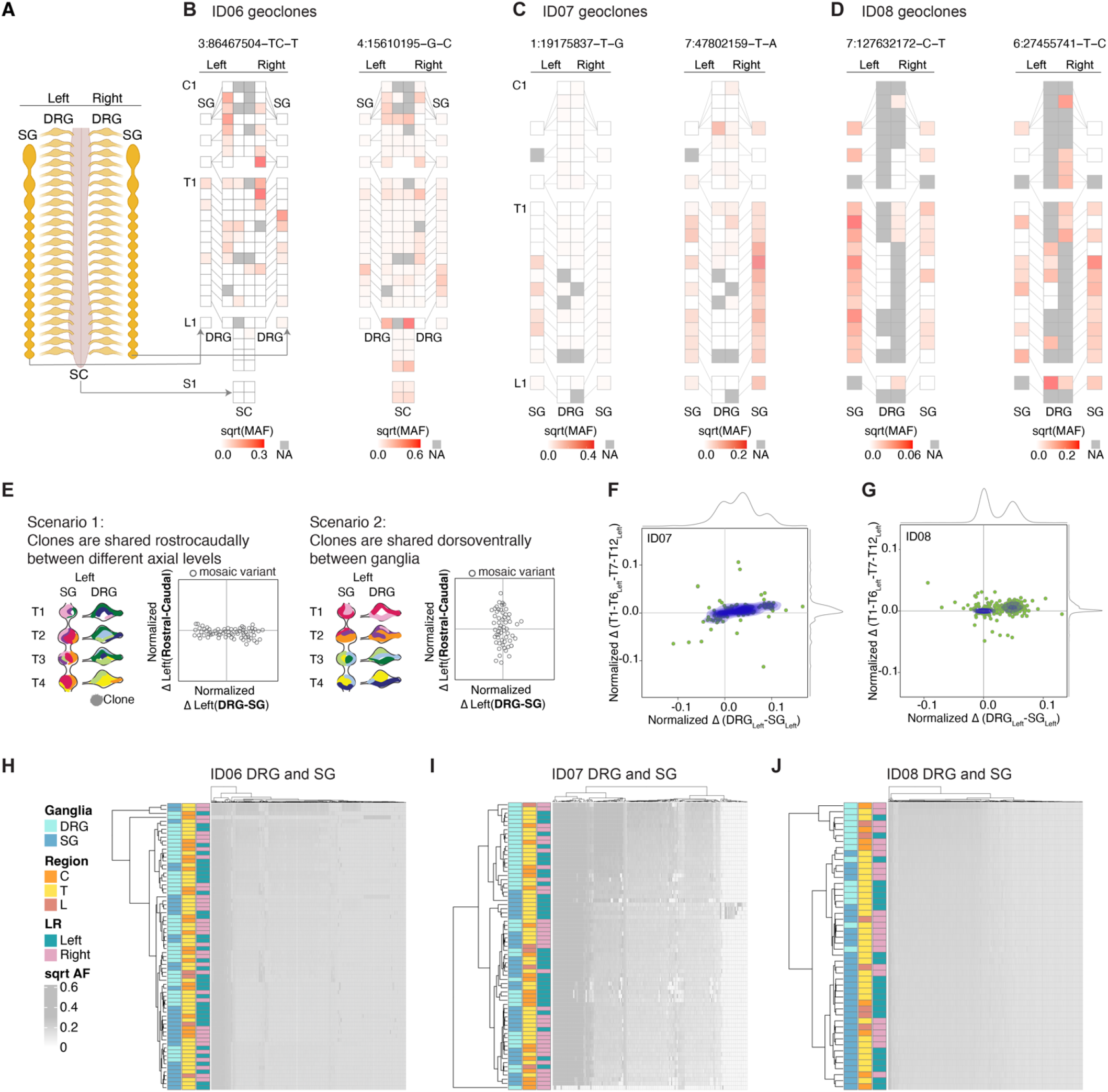
DRGs or SGs share similar lineage relationships along the rostrocaudal axis but are clonally independent. (A-D) Geographical maps of individual MV allelic fractions (AFs) (i.e. ‘geoclones’). Example geoclones in ID06 (B), ID07 (C), and ID08 (D) based upon hg38 reference genomic coordinate (at top). Colors: square root transformed mutant AFs (sqrt-MAF). NA: tissue not available. (E) Models and contour plots of possible scenarios whereby MVs are shared between ganglia along the rostrocaudal axis but restricted dorsoventrally (scenario 1, left) or shared the dorsoventral axis while restricted rostrocaudally (scenario 2, right). Axes: normalized AF difference between rostral and caudal against that between DRG and SG. In scenario 1, most MVs deviate from the center along the x-axis but cluster at the center of the y-axis because of MVs are shared among rostral and caudal levels. In scenario 2, most MVs cluster around the center at the x-axis but not the y-axis since clones are more frequently shared dorsoventrally between DRG and SG, but not between different levels. Clones with genomic similarity are colored similarly. (F-G) MV contour plots observed from ID07 (F) and ID08 (G) thoracic levels with the normalized difference in AFs between rostral (defined as T1-T6 levels) and caudal (defined as T7-T12 levels) (y-axis) plotted against the normalized difference between the DRG and SG (x-axis). Green dots: individual MVs. Blue contours: kernel density estimation of MV distributions. Grey curves: kernel density estimation along the respective axes. Only data for the left-sided ganglia are included in the plots. (H-J) Hierarchical clustering heatmap with Manhattan distances using AFs of MVs from ID06 (H), ID07 (I), or ID08 (J). Note that DRGs and SGs predominantly clustered separately, whereas the left and right tend to intermix together, suggesting the lineage relationship between ganglia is not driven by their anatomical position but rather by their identity (DRG or SG). C: cervical, T: thoracic, L: lumbar.

We wondered if these examples reflected a global clonal structure of DRGs and SGs. By analyzing the total pool of MVs we measured clonal distributions within levels and between levels. We postulated two potential scenarios: 1) a predominant rostrocaudal clonal spread; 2) a predominant dorsoventral clonal spread (i.e. shared between DRG and SG) (Figure 2E). While these two scenarios were not mutually exclusive, we evaluated their prevalence by contour graphs of the normalized difference in the AFs of each MV between the rostral (T1-T6) and caudal (T7-T12) levels against that between the DRGs and SGs (Figure 2E). In extreme cases for the scenarios, rostrocaudal or dorsoventral clonal organization should yield horizontal or vertical contour lines, respectively. Consistent with the genoclone analysis, for most MVs there were substantial AF differences between the DRGs and SGs, with the former primarily quantified with higher AF, presumably reflecting an earlier emergence of DRGs during embryogenesis. Unexpectedly, most ganglia exhibited minimal genomic variation between their rostral and caudal counterparts, forming nearly horizontal contour lines (Figures 2F-2G). We independently performed the analysis for ganglia from the left or right side, yielding similar results (Figures S2M-S2N).

In addition, we compared differences between AFs in ganglia along the dorsoventral and rostrocaudal axes, analyzing the 12 matching pairs of DRG and SG (Figure S2O). To ensure a fair comparison, we used a “rolling levels” approach when quantifying standard deviations along the dorsoventral axis by considering all ganglia from every three spinal levels (see STAR methods). This way, we quantified the individual variance between each of the 12 proximally located ganglia across the rostrocaudal (between spinal levels from T1 to T12) and dorsoventral axis (between DRGs and SGs from every three levels) (Figure S2O). All three donors showed significantly larger genomic variance between ganglia across the dorsoventral than the rostrocaudal axis (Figures S2P-S2R). Together, these findings support a predominant rostrocaudal clonal MVs spread.

### DRGs and SGs show largely genetically distinct lineages

To assess overall lineage relationships of all sampled DRGs and SGs, based on the rationale that ganglia showing the highest clonal similarities should be closest in lineage, we performed hierarchical clustering with Manhattan distances using variant AFs (Figures 2H-2J). In all three donors, prominent left-right lateralized ganglia clusters were absent in the dendrograms. Moreover, there was no evident segregation by spinal levels (cervical, thoracic, or lumbar). However, the DRGs were predominantly clustered away from the SGs. These suggest the lineage relationship of ganglia was not driven by anatomical position or left-right lateralization, but rather by their identity (i.e., DRG or SG).

To establish the robustness of the hierarchical clustering pattern, we performed dendrogram bootstrap analysis (Figures S2S-S2U). Of the 18 significantly stable terminal branches comprised of ganglia with the closest lineage relationship, 17 pairs of ganglia were from 2-6 axial levels apart. This further confirmed the observed rostrocaudal clonal configuration was very unlikely to have occurred by chance.

### Single-nucleus genotyping confirms rostrocaudal clonal structure

Because NC derivatives represented the major cell types of the peripheral ganglia, we hypothesized that they primarily drove the dendrogram structure (Figure S1E). To assess this directly, we performed single-nucleus simultaneous genomic and transcriptomic amplification using the ResolveOME workflow^15,16^, followed by single-nucleus MPAS (snMPAS) and single-nucleus RNA sequencing (Figure 3A, Figures S3A-S3B), profiling a total of 224 single nuclei from the DRGs and SGs of left T2 and T3.

**Figure 3.**
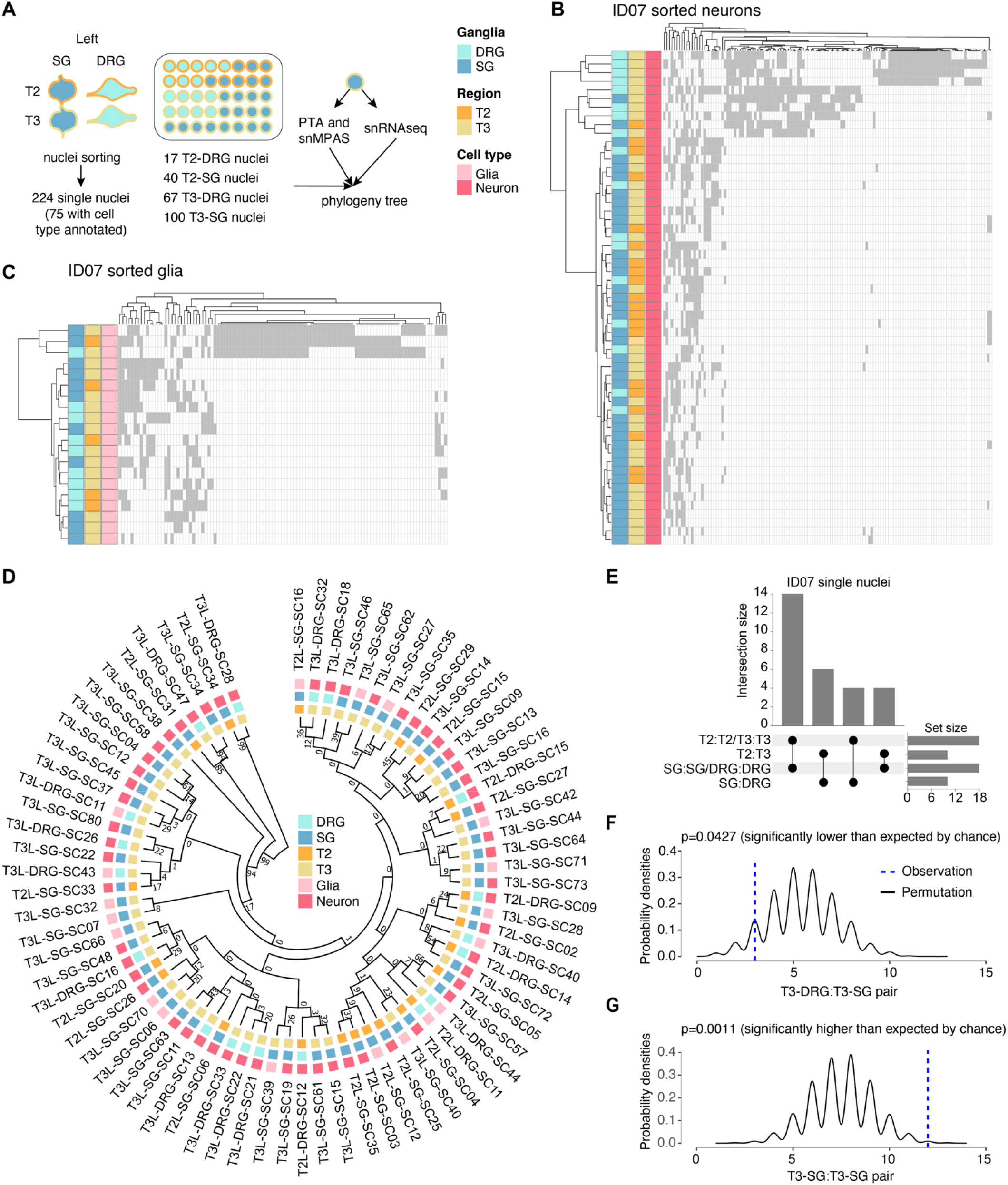
Concurrent single-nucleus MV genotyping with transcriptome analysis confirms the rostrocaudal clonal organization. (A) Strategy for deconvolving the phylogenetic relationship of ganglia at single-cell resolution using ResolveOME (i.e. concurrent DNA amplification by primary template-directed amplification (PTA) and RNA profiling from the same nucleus). MVs genotyped by single-nucleus massive parallel amplicon sequencing (snMPAS) in 224 nuclei isolated from both the DRG and SG at T2 and T3 levels of donor ID07. Of these, cell type was inferred in 75 nuclei by snRNAseq. (B-C) Hierarchical clustering with Manhattan distances from a total of 55 neurons (B) and 20 glia (C) from the DRG and SG at T2 and T3 levels of ID07. Note that DRGs and SGs predominantly clustered separately at both levels, suggesting the lineage relationship between ganglia is not driven by spinal level but rather by identity (DRG or SG). (D) Phylogenic tree following 1,000 bootstrap replications based on the 184 MVs in 75 single nuclei with cell type information. Numbers at branches of the tree: bootstrap values supporting each edge. (E)Upset plot showing number of terminal branches shared between type of ganglia (SG:SG, DRG:DRG or SG:DRG) and axial levels (T2:T2, T3:T3 or T2:T3). (F-G) Number of terminal branches observed in the phylogeny tree in (D) (blue dashed line, observation) and distribution expected after 10,000 permutations (black line: permutation) for T3-DRG:T3-SG pair (p=0.0427, permutation test) (F) and T3-SG:T3-SG pair (p=0.0011, permutation test) (G).

Due to suboptimal RNA quality likely resulting from the post-mortem intervals, we could only confidently annotate the cell type for 75 out of the 224 nuclei using the expression of pan-neuronal or glial marker genes (55 neurons and 20 glia). Nevertheless, hierarchical clustering with the variant alleles in either neurons or glia recapitulated the overall pattern of clonal similarities for DRGs and SGs (Figures 3B-3C). Moreover, to deconvolve lineage relationships among single nuclei, we reconstructed phylogenies for neurons and glia based on snMPAS (Figure 3D). Half of all terminal branches (14/28) were shared by nuclei within a single ganglion, consistent with a predominant local clonal expansion during gangliogenesis (Figure 3E). Remarkably, the remaining branches were mostly comprised of nuclei from different axial levels (10/14), further supporting the rostrocaudal distribution of NC progenitor clones (Figure 3E).

We further performed a permutation test for the phylogeny tree structure by randomly shuffling the labels of the 75 nuclei and computing the probabilities of different terminal branch combinations (Figures 3F-3G). We found that the occurrence of T3-SG:T3-SG pair, indicative of local proliferation in the SG, was significantly more frequent than random (p=0.0011). Notably, the prevalence of the branch shared by T3 DRG and SG was significantly lower than expected by chance (p=0.0427). We further carried out similar analyses for all 224 nuclei and observed similar results (Figures S3C-S3F). This single nucleus analysis suggests that the lineage relationships observed in bulk sequencing are largely driven by neural crest rather than other cell types.

### Determination of neural crest fate precedes specification of left-right axis

The rostrocaudal clonal configuration implied two potential models of trunk NC development: 1) Multipotent NC cells, upon delamination, migrate to prospective DRG and SG, whereupon they are specified to either lineage. Once specified, DRG and SG cells migrate rostrocaudally for a limited range and thereby disseminate clones to neighboring levels; 2) NC progenitors are committed to either DRG or SG prior to delamination, and early rostrocaudal migration of founder cells leads to rostrocaudal clonal organization. In the first model, given that delaminated NC progenitors rarely traverse the midline, likely due to anatomical constraints or extrinsic repellant cues, clones enriched in DRG/SG should primarily lateralize to one side (Figure 4A, left). In the second model, we might expect clones to be predominantly bilateral (Figure 4A, right).

**Figure 4.**
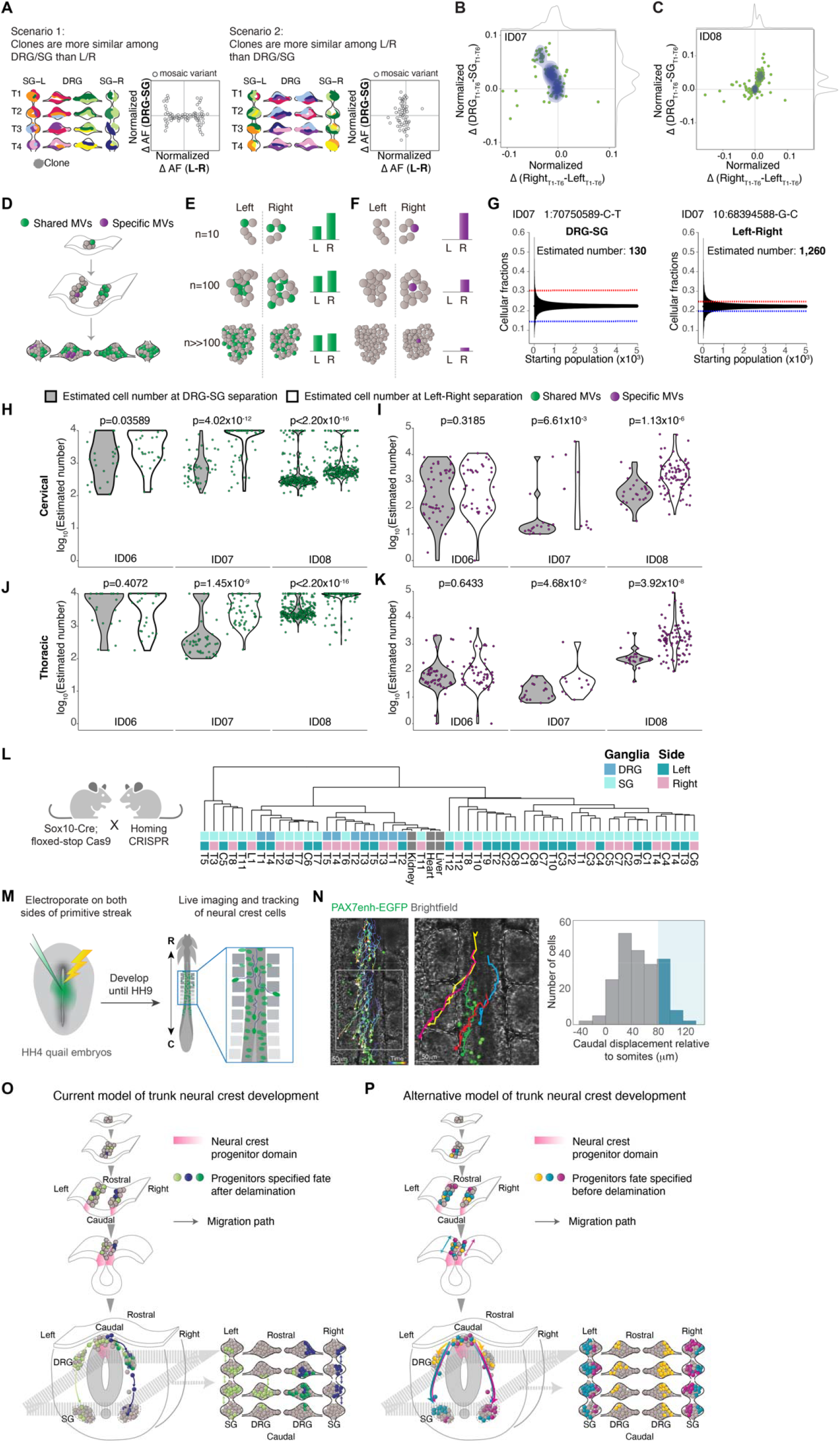
Evolutionarily conserved clonal dynamics and timing of neural crest fate specification. (A) Modeling two possible scenarios whereby clones are more similar among DRG/SG than Left/Right (scenario 1, left) or the opposite case (scenario 2, right). Schematic contour plots showing normalized difference in the allelic fraction (AF) of each MV between the DRG and SG (y-axis) against that between left and right (x-axis). (B-C) Contour plots showing the normalized AF difference of each MV observed from ID07 (B) and ID08 (C) between DRG and SG (y-axis) and between left and right (x-axis). Green dots: individual MVs. Blue contour: 2D Kernel density MV estimation plots highlighted. Grey curves: kernel density estimation along the x-or y-axis. Most MVs spread along the y-axis, indicating a larger difference in AF between DRG and SG, while showing minimal left-right lateral difference. (D-F) Schematics for effect of founder population size MV AFs. Green: MVs acquired during early embryogenesis before left-right split. Green variants therefore are shared in ganglia of both left and right but with varying AFs, whereas purple MVs are acquired only after the left-right split, thus distributed on one side exclusively. (E) AFs for green variant quantified under three hypothetical founder population cell numbers: n=10, n=100, or n>>100 shown. Larger hypothetical founder population size correlated with smaller AF difference between left and right. (F) For the purple lateralized variant, smaller number of cells immediately after left-right lateralization correlated with higher AFs. (G) Estimation of the effective population size by the observed difference in AF as in (D-F). Representative variants from ID07 for calculating the cell number prior to DRG-SG split (left panel) and prior to left-right split (right panel). Blue and red dashed lines: difference in average AF for the individual variant between DRG and SG (left panel) or between left and right (right panel). Black lines: 95% bands of hypergeometric distribution from each simulated starting population size. (H-K) Violin plots comparing estimated maximum number of founder population size at DRG-SG specification and left-right lateralization in cervical regions (H) or thoracic regions (J). Green dots: MVs distributed bilaterally; Violin plots comparing the estimated minimum size of the founder population at DRG-SG specification and left-right lateralization in cervical (I) or thoracic regions (K). Purple dots: MVs restricted to one side. The predicted population size at DRG-SG specification is significantly smaller than at left-right split when estimated from either class of MV. P-values: two-tailed Mann-Whitney U tests. (L) Mouse breeding scheme for generating the neural crest-specific CRISPR barcoding mouse model. Migratory neural crest-specific *Sox10-Cre* driven Cas9 activity for *in vivo* barcode editing. Representative lineage tree dendrogram for the edited barcodes from 1-month-old mice. Bulk organs (liver, kidney, and heart) with limited NC contribution (n=2 independent mice). (M)Experimental design for real-time *ex ovo* imaging of NC progenitor migration in quail embryos. (N)Representative tracks showing migration paths of NC cells expressing H2B-Citrine under control of *Pax7* enhancer. Histogram showing the rostrocaudal migration distance of 212 cells tracked (n=6 embryos). Blue shading indicates cells migrating caudally by more than 1 somite (mean somite length 81 ± 1 µm, n=92 somites from 6 embryos). Right panel: magnification of boxed region from left panel. Brackets: somites. Scale bar: 50μm. (O-P) Trunk NC development current model (O) and the alternative model observed from this study (P). The current model has little in the way of rostrocaudal cell movement prior to NT closure, and that individual clones populate both DRG and SG (depicted by green or purple cells in both types of ganglia, mostly restricted to one side). The alternative model includes rostrocaudal cell movement prior to NT closure, and that individual clones populate either DRG or SG bilaterally but infrequently populate both DRG and SG (depicted by yellow cells in DRGs on both left and right, and both turquoise and violet cells in SGs on both left and right).

Therefore, we compared the AF difference for each MV between the left and right ganglia against that between the DRGs and SGs in corresponding axial levels. Intriguingly, most MVs exhibited fewer AF differences between left and right while simultaneously showing higher AF variation between DRGs and SGs (Figures 4B-4C and S4A-S4B). Accordingly, bilateral clones enriched in either type of ganglia favored model 2. We next tested whether NC progenitors are fate committed before delamination from dorsal NT, i.e. the theoretical latest timepoint for left-right commitment. We estimated the relative population sizes at these two developmental events, by computational simulation. For any MV shared between DRG-SG or left-right, the smaller the AF difference between groups (left vs right or DRG vs SG), the larger the estimated population size was at the time the group segregated (Figures 4D-4E, STAR Methods). We thus performed a stepwise simulation to estimate the maximum possible cell number when NC progenitor fate specification or left-right commitment likely occurred (Figure 4G).

By contrast to MVs that were shared between groups, the observed AF for MVs fully lateralized or exclusive in either ganglion predicted the lower limit of cell number immediately following lateralization or cell fate specification respectively, assuming the MV is nascently acquired in one progenitor (Figures 4D and 4F). Although the simulated population size difference for donor ID06 was modest, in part due to AF distribution likely skewed by local clonal proliferation, both the upper and lower limit of estimated cell number at NC progenitor fate specification was substantially smaller than at left-right separation for both donors ID07 and ID08 (Figures 4H-4I). Notably, the smaller estimated cell number was consistent across cervical and thoracic levels (Figures 4J-4K), suggesting NC fate specification occurred at an earlier developmental time point. These findings support NC fate specification preceding dorsal NT delamination.

### NC clonal organization is evolutionarily conserved

To confirm whether the relative clonal independence between DRGs and SGs is evolutionarily conserved, we utilized the Homing CRISPR barcoding mouse line^17^ crossed with *Sox10-Cre* to drive NC-specific Cas9-induced recombination, then microdissected individual DRG and SGs to generate datasets comparable to those in human. *Sox10* is first expressed by NC cells as they migrate from the NT and thus Cas9 editing should be limited to cells dividing after delamination^18^. The lineage tree of DRGs and SGs in mice was then reconstructed based on the percentage of cells sharing identical edits across multiple MVs. Similar to the clonal organization observed in humans, we found that the DRGs across multiple axial levels displayed more clonal similarities than DRGs and SGs within similar levels (Figures 4L). Moreover, as observed in humans, the SGs and DRGs clustered away from each other, consistent with independent lineage relationships (Figure S4C). Next, to directly observe NC progenitor dynamics, we performed real-time imaging of quail embryos electroporated with the early NC-specific *Pax7* enhancer reporter (Figure 4M). *Pax7* is expressed in cells fated to become NC at the neural plate border^9^. Tracking of cell migratory paths revealed more than 20% of NC progenitors migrate along the rostrocaudal axis across at least two somite levels within 6 hours of imaging (Figure 4N and Movie S1). To assess whether the bilateral clones are contributed by NC cells migrating across the midline prior to delamination, we electroporated *FoxD3* reporter to one side of the quail embryo NT. *FoxD3* expresses later than *Pax7* but is one of the earliest premigratory NC markers^19^. Notably, we observed considerable migration of *FoxD3-*positive cells across the midline to take up contralateral positions prior to NT delamination (Figures S4D-S4E). These observations in rodents and avians support that the bilateral, rostrocaudal clonal dynamics of trunk NC are evolutionarily conserved.

## Discussion

Here we perform a comprehensive, large-scale analysis of trunk NC clonal dynamics and uncover lineage relationships between the major cellular derivatives. Our results suggest unexpected developmental organization of the NC cells, which may require a revision of current models of NC cell fate diversification, with relevance to the origins and pathology of the broad spectrum of clinical ‘neurocristopathies’.

### Rostrocaudal NC migration before delamination distributes progenitor clones bilaterally

It is generally perceived that trunk NC cells mostly migrate in a multimeric, segmented organization^20^, with cells migrating exclusively through the rostral but not the caudal half of the sclerotome. Only after cells have reached the ventral edge of sclerotome in the vicinity of dorsal aorta, do they extend interganglionic filopodia and form a contiguous narrow stream spanning two segments rostrally or caudally from their axial origins^21^. These filipodia-contact-based cell intermingling should result primarily in clones shared between contiguously located SGs. Therefore, rostrocaudal migration was thought to be possible only within a narrow temporal and spatial window, primarily reserved for cells colonizing the SG, as the cells destined for DRG were not expected to migrate ventrally past the sclerotome. Corroborating this model, previous studies with chick-quail transplantation showed that NC cells from a single somite level can contribute to SG spanning several axial levels but generally only colonize the DRG at the corresponding segment^21,22^ (Figure 4O).

However, this model fails to explain several of our observations: 1) We found numerous clones populating DRGs separated by multiple axial levels; 2) We found existence of multiple bilateral clones specific to either DRG or SG; 3) We found clones not shared between adjacent levels but rather disseminated across the entire thoracic region. Thus, we think the prior model may require some revision.

Here our data from human and mouse compelled us to live imaging in quail embryos, which confirmed robust NC cell migration along the rostrocaudal axis prior to their delamination. These cellular movements disseminated NC progenitor clones across different axial levels, driving the rostrocaudally shared clonal organization. Meanwhile, a substantial proportion of NC progenitors crossed the midline prior to their delamination. Together these migratory behaviors potentially explain the bivalent, multiple levels-spanning distribution of NC progenitor clones (Figure 4P). These observations also contrast with differences between the central and peripheral nervous systems, where in the brain we and others showed that the midline axis is established initially, and that progenitors strictly respect the midline^14,23^.

### “Early” fate restriction is the prevalent mode of NC development in mammals

The pivotal question of whether NC progenitors are multipotent has been debated for decades. Conflicting evidence supports two possible models: one posits that the entire NC population is multipotent, capable of generating multiple cell types in response to environmental cues^6,24-26^; the other suggests that the NC progenitor pool is heterogeneous, comprising spatiotemporally restricted cells with fate predetermined before NT delamination^27-29^.

Vital dye or viral fate mapping studies have the potential to yield conflicting results, in part due to the caveat of inadvertently labeling multiple rapidly expanding mitotic cells. The recent study using *Confetti* multicolor fluorescent reporter transgenic mice was thought to settle the longstanding controversy, suggesting NC retained multipotency after delamination^6^ (Figure 4O). However, analysis was limited to a single axial level, and thus may have missed potential rostrocaudal movement. Rostrocaudal clonal spread demonstrated here suggests the intriguing possibility that identical combinations of fluorescent proteins in a single transverse section may not necessarily derived from the same clone, but rather represent distinct clonal origins from different axial origins.

Through mathematical simulation of the population size at the timing of cell fate specification and left-right commitment, we suggest that in most cases the restriction of NC progenitor fate precedes left-right commitment. These results also suggest lineage commitment to either DRG or SG may occur while cells are still NT-resident. However, while this represents the *in vivo* scenario, we cannot rule out the possibility that NC possesses inherent plasticity to change fate *in vitro*. Also, our single-cell clonal analysis suggests that a few cells from the DRG and SG at the same level may share the closest lineage, although this occurrence is less frequent than expected by chance. Therefore, our findings also confirm the presence of multipotent NC cells, but likely constituting only a minor population (Figure 4P).

### Mulitmodal assessment of embryonic cellular origins in vertebrates

Compared with conventional lineage tracing methods, MVBA offers several significant advantages: 1) MVs imprinted in the genome allow faithful recapitulation of lineage history; 2) No genetic or surgical manipulation is required, allowing for assessment in human; 3) Because the entire genome is assessed, the probability of two unrelated cells sharing an identical variant by chance is extremely low. MVBA in human also offers potential advantages over lineage tracing using in vivo or in vitro models: 1) While lineage tracing in genetic models like CARLIN or *Confetti* mice is attractive, it is not possible to exclude that the same genetic variant arose twice independently^30,31^; 2) MVBA circumvents the inevitable caveat of reactivating stemness due to in vitro culture microenvironment, which could produce false-positive lineage relationships^32^; 3) MVBA allows assessment of lineage relationships between anatomically distant tissues, which may be difficult with Cre-driven recombination. The major limitation of MVBA in human is the cost and ethical considerations, as well as inability to directly assess timepoints or visualize cellular dynamics. To overcome these limitations, we performed real-time imaging of whole avian embryos to directly observe progenitor migratory behavior, revealing previously unappreciated robust trunk NC cellular movements. Therefore, our study combines several contemporary lineage tracing methods to bring insights into a classical biological question.

### Limitations of the study

It is ethically impossible to perform live imaging of human post-gastrulation embryos, so instead we inferred cell lineage from naturally occurring MVs on a limited scale. Thus, while our findings suggest that some DRG and SG lineages may be determined before cells exit the NT, in contrast to prevailing models, we were not able to directly assess cell movement in the human fetus. Because of the small tissue size and relatively limited cell number in each ganglion, it was technically challenging to isolate individual cell types for genomic analysis. To overcome this hurdle, we performed single-nucleus ResolveOME analysis from a subpopulation of cells. However, large-scale lineage deconvolution on the cell-type level remains unpractical. Due to the postmortem interval of human cadavers, single-nucleus transcriptional profiling was only possible in a minority of nuclei assessed.

## Acknowledgments

We thank all the individuals who donate their bodies and tissues for the advancement of scientific research. We thank Dr. Marianne Bronner (Caltech) and Dr. Ruth Williams (U Manchester) for the generous gift of the FoxD3enh-GFP and Pax7enh-GFP plasmids, respectively, Cody Fine, Mateo Espinoza and Mitra Banihassan (UC San Diego) for FACS technical support, UCSD Stem Cell Program and CIRM Major Facilities grant (FA1-00607) at the Sanford Consortium for Regenerative Medicine, NIMH (grants U01MH108898 and R01MH124890 to J.G.G. and R21MH134401 to X.Y.), the Larry L. Hillbolm Foundation (to J.G.G.), NICHD (grants P01HD104436 to J.G.G. and K99/R00HD111686 to X.Y.) and the Rady Children’s Institute for Genomic Medicine, the San Diego Supercomputer Center (SDSC) (TG-IBN190021 to X.Y. and J.G.G.), and SIG grant (S10OD026929) to UCSD IGM, S10OD021644 supporting the Center for High Performance Computing (CHPC) and Utah Center for Genetic Discovery (UCGD) at the U Utah.

## Author Contributions

K.I.V., X.Y. and J.G.G. conceived the project and designed the experiments. K.I.V., X.Y., C.C., R.H., N.M., Q.Z., C.B., A.P., N.C., S.F.K. performed experiments and data analysis. Y.D.A., J.X.W., and M.D.W. performed live imaging and data analysis. G.N. and S.T.B. performed dissection of human donors; K.I.V. and F.J. performed mouse dissections; K.I.V., X.Y. and J.G.G. drafted the manuscript with input from all authors.

