## Supplemental Information for "Genomic mosaicism reveals developmental organization of trunk neural crest-derived ganglia"

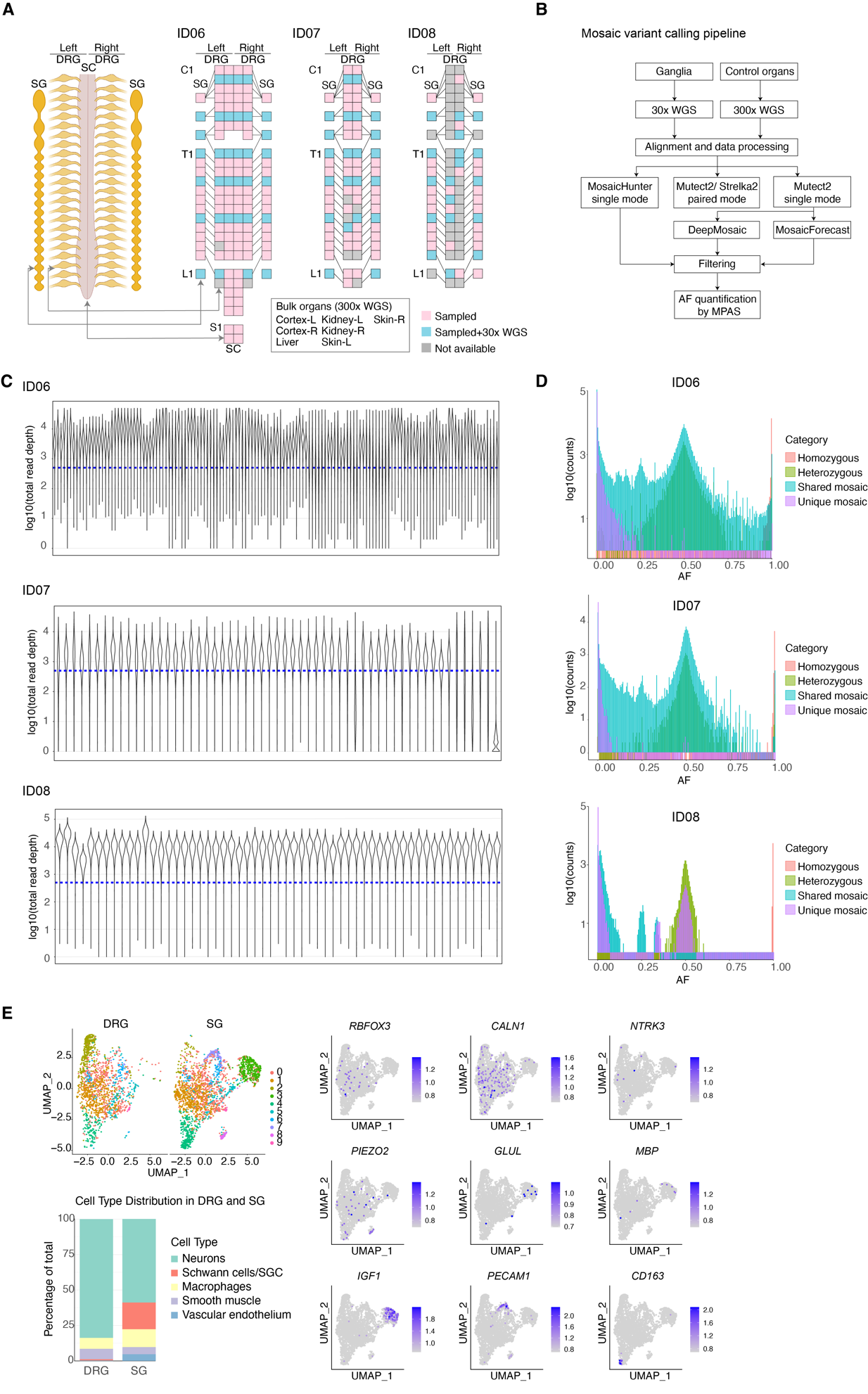

Figure S1. Tissue collection from human donors and quality controls of MPAS, related to Figure 1.

(A) Schematic showing the collection of dorsal root ganglia (DRG), sympathetic ganglia (SG), and other major organs from the 3 adult donors. Pink boxes: ganglia that were sampled; blue boxes: ganglia processed for 30x whole genome sequencing (WGS); grey boxes: not analyzed due to suboptimal tissue integrity or sequencing quality. C, cervical; T, thoracic, L, lumbar; S, sacral; SC, spinal cord.

(B) Bioinformatic workflow for the identification of candidate mosaic variants. See STAR Methods for a detailed description.

(C) Violin plots of log-transformed total read depths (y-axes) of all somatic mosaic variants in 140 samples from ID06 and 57 from either ID07 or ID08 (x-axes). The blue dashed lines indicate 500x read depth.

(D) The distribution of allelic fraction of mosaic variants categorized into homozygous, heterozygous, shared mosaic, or unique mosaic variants in all the samples from the 3 donors. AF, allelic fraction.

(E) Estimation of cell type proportion in the DRG and SG from donor ID08 by single-nucleus RNA sequencing. Uniform manifold approximation and projection (UMAP) graph from single-nucleus RNA sequencing with nuclei isolated from DRG (n=1,714) and SG (n = 2,652) (left upper panel). Feature plots for expression of known markers of major cell types in the ganglia (right panel). The identity of cell clusters was annotated with marker expression. Bar graph showing cell type proportion in the DRG and SG (left lower panel).

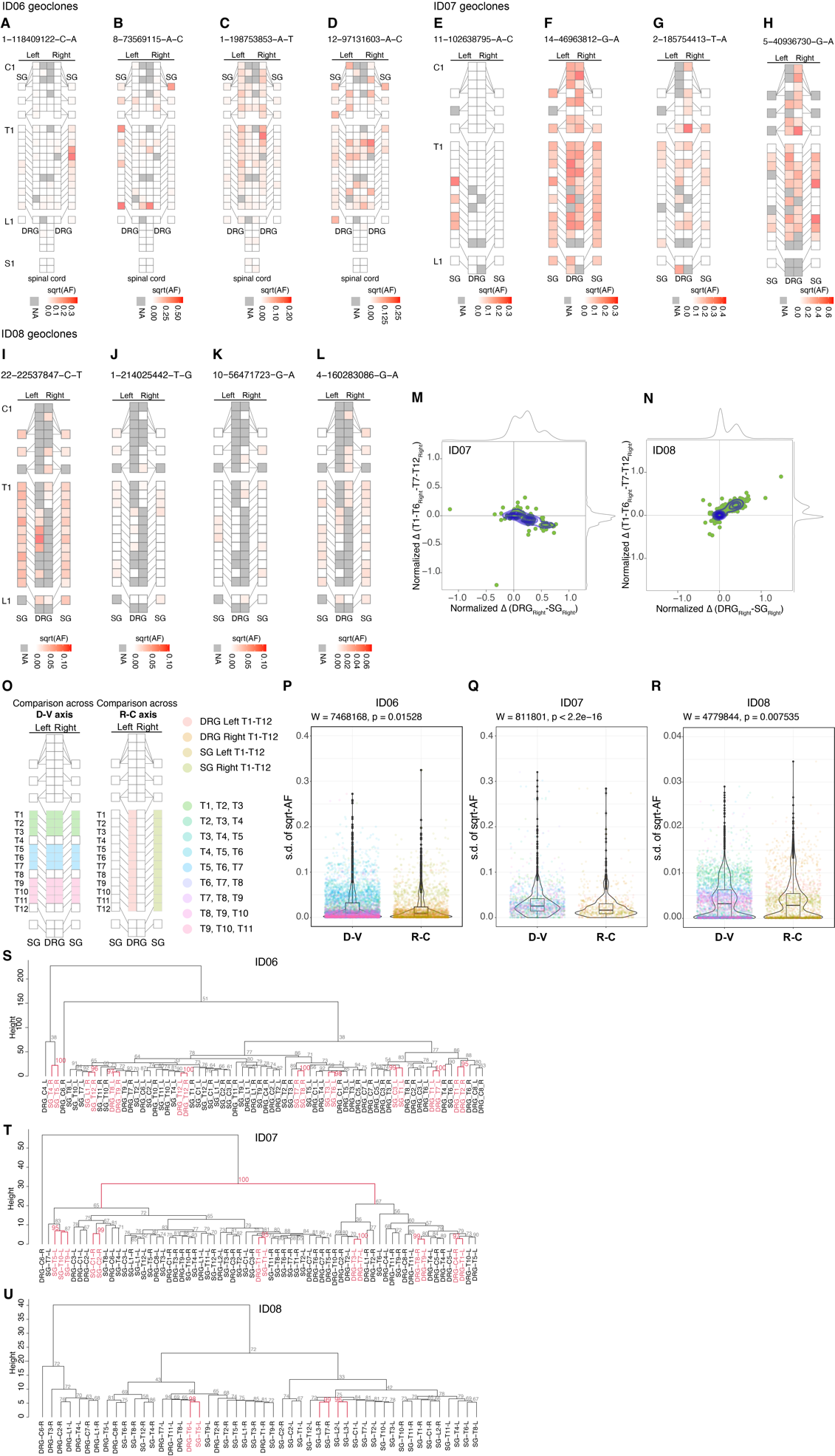

Figure S2. Clonal dynamics of the human DRG and SG, related to Figure 2.

(A-L) Geoclone representation of the square-root transformed allelic fraction of example variants from ID06 (A-D), ID07 (E-H) and ID08 (I-L).

(M-N) Contour plots of the normalized difference in the allelic fractions between rostral (defined as T1-T6 levels) and caudal (defined as T7-T12 levels) of the right side (y-axis) against the normalized difference between the DRG and SG of the right (x-axis) for each mosaic variant from ID07 (M) or ID08 (N). Each dot represents an individual mosaic variant. Kernel density estimation of the mosaic variant distributions highlighted in purple. Grey curves represent the kernel density estimation along the respective axes.

(N) Contour plots of normalized difference in the allelic fraction of each mosaic variant observed from ID07 between DRG and SG (y-axis) and that between left and right (x-axis) for samples between T7 and T12 levels. Each dot represents an individual mosaic variant. 2D Kernel density estimation plots of mosaic variants highlighted in purple. Grey curves are the kernel density estimation along the x- or y-axis.

(O) Schematics showing the comparison in the difference of variant allelic fractions along the dorsoventral (D-V) axis (between DRG and SG) and along the rostrocaudal (R-C) axis (between all thoracic levels).

(P-R) Violin plots showing the standard deviation of square-root transformed allelic fraction (s.d. of sqrt-AF) for the variants from ID06 (P), ID07 (Q), and ID08 (R) compared between the D-V axis and R-C axis. Each dot represents individual mosaic variants, with the color of the dots indicating the specific ganglia from which standard deviation was quantified, either along the D-V or R-C axis. P values are from two-tailed Mann-Whitney U tests. W statistics are the sum of the ranks.

(S-U) Bootstrapping results of the lineage dendrogram for ID06 (S), ID07 (T) and ID08 (U). The numbers next to each branch represent the bootstrap values after 10,000 replicates. Red-colored branches indicate the samples with approximated unbiased p-value > 95%, rejecting the hypothesis “the cluster does not exist” at a statistically significant level (<5%).

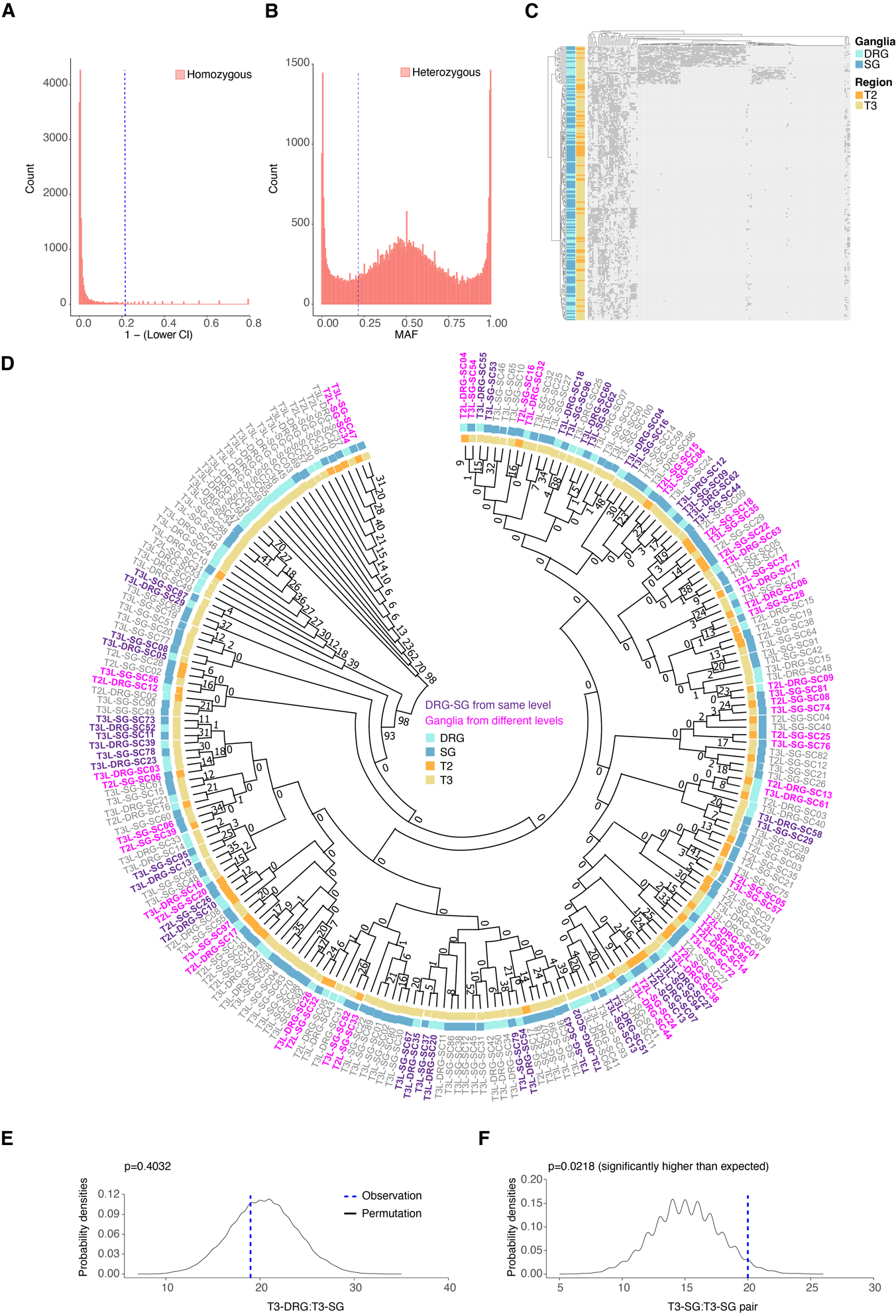

Figure S3. Single-nucleus genotyping of mosaic variants in DRG and SG from T2 and T3 levels, related to Figure 3.

(A) False positive rate of mosaic variant genotyping by snMPAS. Distribution of the reference homozygous variants from ID07, with the x-axis representing 1 minus the lower bound of 95% binominal confidence interval (95% CI). The blue dashed line indicates the cut-off for a false discovery rate of 5%.

(B) False negative rate of mosaic variant genotyping by snMPAS. Distribution of the upper bound of 95% binomial confidence interval. The blue dashed line indicates the cut-off for a false discovery rate of 5%

(C) Heatmap of the hierarchical clustering with Manhattan distances of the allelic fractions of mosaic variants in all 224 nuclei that underwent snMPAS following primary template-directed amplification (PTA).

(D) Phylogenic tree after 1,000 bootstrap replications from the 184 MVs in the 224 single nuclei. The numbers at branches of the tree are bootstrap values supporting each edge. Purple indicates a terminal branch pair that consists of both DRG and SG from the same level (either T2:T2 or T3:T3). Pink indicates ganglia pairs from different levels at the terminal branch (T2:T3).

(E-F) Graphs comparing the actual number of terminal branches observed in the phylogeny tree (blue dashed line, observation) and the distribution of expected number after 10,000 permutations (black line, permutation) for T3-DRG:T3-SG pair (p=0.4032, permutation test) (E) and T3-SG:T3-SG pair (p=0.0218, permutation test) (F).

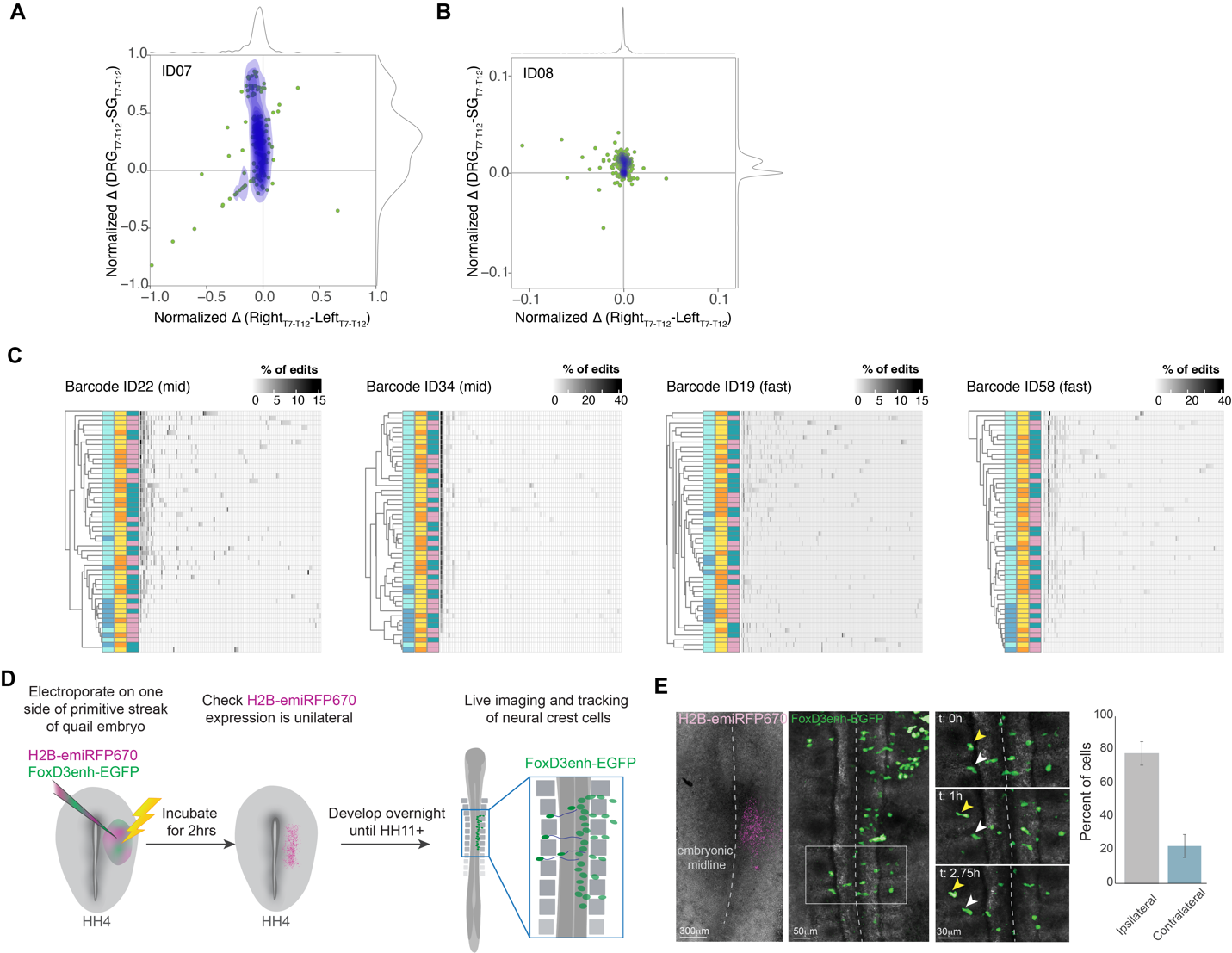

Figure S4. Evolutionarily conserved clonal dynamics of trunk neural crest, related to Figure 4.

(A-B) Contour plots showing the normalized difference in the allelic fraction of each mosaic variant observed from ID07 (A) and ID08 (B) between DRG and SG at T7-T12 levels (y-axis) and between left and right (x-axis). Green dots: individual mosaic variant. Blue contour: 2D Kernel density estimation. Grey curves: kernel density estimation along the x- or y-axis

(C) Representative hierarchical clustering of CRISPR/Cas9 edits at individual barcode locus.

(D-E) Schematic of experimental design to evaluate midline-crossing migration of NC progenitors. Quail embryos at Hamburger-Hamilton stage 4 (HH4) were electroporated with H2B-RFP and *FoxD3* enhancer driven EGFP on one side. Spatial specificity of electroporation was confirmed by inspecting H2B-RFP expression 2 hr later. Live imaging and tracking of FoxD3+ NC progenitors were then performed at HH11 (D). Quantification of cells migrating on the side of electroporation (ipsilateral) and crossing the midline (contralateral) indicated 20% of NC migrated to the opposite side during the 6 hours of imaging (n=5 embryos) (E).

Movie S1. Live imaging of rostrocaudal migration of early NC progenitor cells in the quail embryo, related to Figure 4.

Cells expressing H2B-citrine under the control of the *Pax7* enhancer migrate along the rostrocaudal axis during neural tube closure in the quail embryo. The movie shows all labelled cells initially (00:00-00:06) and then repeats, showing all cells tracked (00:06-00:12), then the subset of tracked cells shown in Figure 4N (00:12-00:24) and finally the position of those tracks relative to neighboring somites (00:25-00:29, shaded in grey).
